# G protein activation occurs via a largely universal mechanism

**DOI:** 10.1101/2023.10.24.563885

**Authors:** Neha Vithani, Tyson D. Todd, Sukrit Singh, Kendall J. Blumer, Gregory R. Bowman

**Affiliations:** Department of Biochemistry and Molecular Biophysics, Washington University School of Medicine, St. Louis, Missouri 63110, United States; Center for the Science and Engineering of Living Systems (CSELS), Washington University in St. Louis, St. Louis, Missouri 63130, United States; Department of Cell Biology and Physiology, Washington University School of Medicine, St. Louis, Missouri 63110, United States; Departments of Biochemistry & Biophysics and Bioengineering, University of Pennsylvania, Philadelphia, PA 19104, United States

## Abstract

Understanding how signaling proteins like G proteins are allosterically activated is a long-standing challenge with significant biological and medical implications. Because it is difficult to directly observe such dynamic processes, much of our understanding is based on inferences from a limited number of static snapshots of relevant protein structures, mutagenesis data, and patterns of sequence conservation. Here, we use computer simulations to directly interrogate allosteric coupling in six G protein α-subunit isoforms covering all four G protein families. To analyze this data, we introduce automated methods for inferring allosteric networks from simulation data and assessing how allostery is conserved or diverged among related protein isoforms. We find that the allosteric networks in these six G protein α-subunits are largely conserved and consist of two pathways, which we call pathway-I and pathway-II. This analysis predicts that pathway-I is generally dominant over the second pathway, which we experimentally confirmed by showing that mutations to pathway-I perturb nucleotide exchange more than mutations to pathway-II. In the future, insights into unique elements of each G protein family could inform the design of isoform-specific drugs. More broadly, our tools should also be useful for studying allostery in other proteins and assessing the extent to which this allostery is conserved in related proteins.

## Introduction

Understanding how signaling proteins like G proteins are allosterically activated is a long-standing challenge with significant biological and medical implications. Insight into G protein activation is of great biological significance because these proteins play key roles in regulating a wide range of biological processes including vision, olfaction, taste, neurotransmission, cell proliferation, and motility (Johnston & Siderovski, 2007; Oldham & Hamm, 2006). These processes are mediated by 16 Gα protein isoforms, which are categorized into four families based on downstream signaling modality and sequence homology: Gα_i/o_, Gα_s_, Gα_q/11_, and Gα_12/13_ (Anantharaman et al., 2011; Strathmann & Simon, 1991). Dysfunction of these different isoforms results in diseases ranging from cancer to pseudohypoparathyroidism (Lania et al., 2006; Schöneberg & Liebscher, 2021; Spiegel & Weinstein, 2004). One strategy to treat these diseases is to target the relevant G protein isoform without eliciting off-target effects by simultaneously impacting other isoforms. Achieving this end would be facilitated by understanding how each isoform is activated and identifying structural elements that are shared or unique to each isoform. Such structural elements could be unique either because they play a more important role for some isoforms than others or because they are important in multiple isoforms but have unique sequences/chemistry in some. Ideally, these subtle differences could then be utilized to yield therapeutics with the desired specificity and minimize off-target impacts by uniquely targeting shared, but sequence diverged, structural elements or structural elements with disparate isoform-specific effects.

Interrogating dynamic processes like the allosteric coupling responsible for G protein activation remains difficult despite significant progress in understanding the structure and function of these molecular switches. G proteins are heterotrimeric complexes composed of an α, β, and γ subunit (Hurowitz et al., 2000). The α subunit (abbreviated Gα and shown in Fig. 1) is the key switch, which in the inactive state has GDP bound between its two domains, the ras-like G domain and the alpha-helical H domain. G protein-coupled receptors (GPCRs) activate G proteins by binding the Gα subunit and triggering exchange of GDP for GTP with subsequent separation from Gβγ to yield the active form of Gα. However, many questions remain about how the GPCR- and nucleotide-binding sites communicate given that they are over 30 Å apart. Analysis of sequence conservation and a wealth of biochemical data suggested all G proteins use a universal activation mechanism (Flock et al., 2015). However, it is necessary to test this hypothesis by directly observing the activation process of every G protein subtype. Molecular dynamics simulations provide a means to study with atomistic detail how similar the allosteric networks responsible for G protein activation are across families, but so far publications on this subject have focused on single G proteins (X. Sun et al., 2018; Yao et al., 2016).

**Figure 1.**
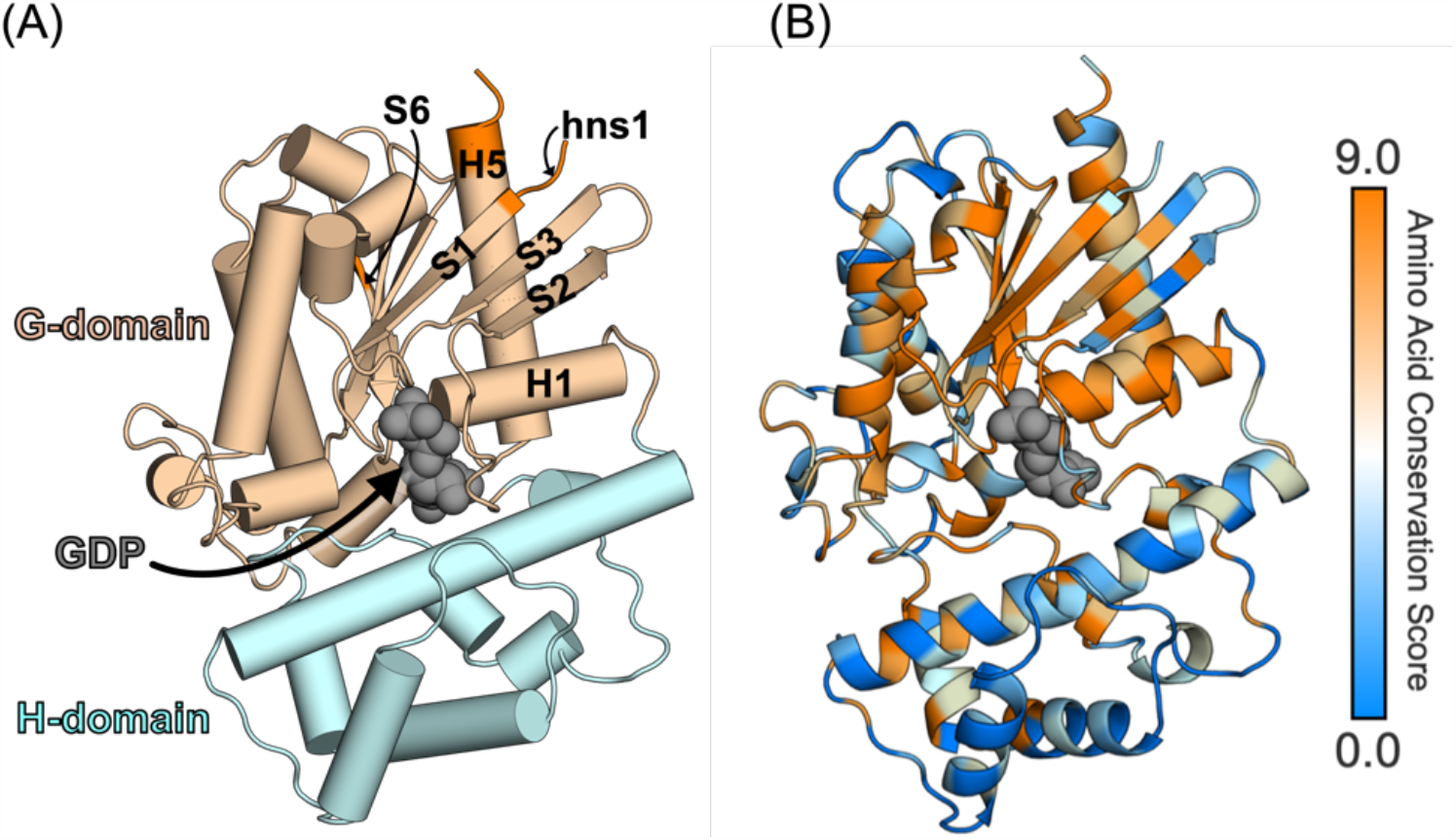
Structure of a Gα subunit and sequence variation across the 16 Gα isoforms. (A) Structure of Gq (PDB: 3ah8) highlighting GDP (grey spheres) bound to the active site, which is between the G domain (light brown) and H domain (cyan). Key secondary structure elements are labeled according to the CGN system. GPCR-binding regions are shown in orange (B) Amino-acid conservation score computed on the Consurf server using the multiple sequence alignment of the 16 G*α* isoforms mapped onto the structure of Gq.

Here, we use molecular dynamics simulations to assess whether the four G protein families use a universal allosteric network to become activated. Specifically, we ran an aggregate of over 210 μs of simulation of six different Gα isoforms: Gi1, Gt, and Go from the Gi/o family; Gq from the Gq/11 family; G13 from the G12/13 family; and Gs from the Gs family. For each isoform, we ran tens of μs of molecular dynamics simulations, built a Markov state model (MSM) to represent the conformational space explored, and identified the allosteric networks connecting the GPCR- and nucleotide-binding sites, as we have done previously for Gq (X. Sun et al., 2018). We also present new approaches for assessing the conservation of allosteric networks. We focus on assessing the extent to which the different G proteins share a universal activation mechanism. To discuss comparisons between isoforms, we use the Common G*α* numbering (CGN) system (Flock et al., 2015). In this system, residues are named based on their position in a secondary structure element. For example, Phe^G.H5.8^ refers to the Phe at the 8^th^ residue of the fifth helix of the G domain and Cys^G.s6h5.2^ refers to the Cys at the 2^nd^ residue of the loop between β strand 6 and helix 5 of the G domain. In the future, drawing on our work to delve deeper into the differences between subfamilies/isoforms will likely be of great value for informing the development of inhibitors that specifically target different subsets of G proteins.

## Methods

### Molecular dynamics simulations of Gα isoforms

Crystal structures of Gαi1 (PDB id: 3QE0) (Bosch et al., 2012), Gαt (PDB id: 1GOT) (Lambright et al., 1996), Gαo (PDB id: 3C7K) (Slep et al., 2008), Gα13 (PDB id: 1ZCB) (Kreutz et al., 2006) and Gαs (PDB id: 6EG8) (Liu et al., 2019) were used to set up simulations of GDP-Gα for these isoforms. A Mg^2+^ ion coordinating to GDP at the active site was also including in each system. Helix HN at the N-terminal and residues after position G.H5.21 at the C-terminal were removed from these structures to make the protein lengths uniform across isoforms including Gαq system prepared in our previous study (Sun et al., 2018).

Each protein structure was solvated in a dodecahedron box of TIP3P water model (Jorgensen et al., 1983) that extended 10 Å from the protein in every dimension. Na^+^ and Cl^-^ were added to prepare a neutral system with 0.15 M NaCl. AMBER03 force field was used for protein (Duan et al., 2003). The force field parameters of GDP were obtained from the AMBER Parameter Database (Meagher et al., 2003) as described in our earlier work on Gαq (Sun et al., 2018). Virtual sites were used to allow for a 4 fs time-step (Feenstra et al., 1999) during the production runs of molecular dynamics.

Solvated protein systems were energy minimized using steepest descent algorithm for 50000 steps until the maximum force reached below 100 kJ/mol/nm. A step size of 0.02 nm and a cut-off distance of 0.9 nm for the neighbor list, electrostatic interactions and van der Waals interactions were used during the energy minimization step. Subsequently, NVT simulations were run with the constraint of 1000 kJ/mol/nm applied to the protein heavy atoms using a time step of 2 fs. During these simulations, the temperature of the system was constrained to 300 K using V-rescale thermostat with a time constant of 0.1 ps (Bussi et al., 2007). A cut-off distance of 1 nm was used for the van der Waals, short-range electrostatic interactions and periodic boundary conditions were applied to x, y and z directions. The Particle-Mesh-Ewald method was employed to recover the long-range electrostatic interactions with 0.16 nm as the grid spacing and with a fourth order spline (Kolafa & Perram, 1992). All the bonds connecting to hydrogens are constrained using the LINCS algorithm (Hess, 2008). Following the NVT simulations, the systems were equilibrated using NPT simulations for 1 ns using the integration time step of 2 fs. All simulation parameters were the same as those used during NVT equilibration. In addition, Parrinello-Rahman barostat for pressure coupling (Parrinello & Rahman, 1981) was applied to constrain the pressure of the system to 1 bar. The position constraints on heavy atoms were removed during the subsequent production runs set up on Folding@home. We performed 100 parallel simulations of the GDP-bound state of Gα isoforms on the Folding@home (Shirts & Pande, 2000) distributed computing platform. For each isoform, aggregate simulations of more than 35 microseconds were performed.

### Correlation of All Rotameric and Dynamical States (CARDS) analysis

We applied the CARDS (Sun et al., 2018) framework to derive inter-residue structural and dynamical correlations to understand allosteric communications in Gα isoforms (Fig. S1). The CARDS method measures correlations between every pair of dihedral angles (backbone and side chain) by quantifying correlated changes in their rotameric (structural) states and dynamical behavior. Structural states of backbone dihedrals are determined by categorizing Φ and Ψ dihedrals into rotameric states cis and trans, respectively. The side-chain dihedrals are categorized in three states gauche+, gauche-, and trans. Every dihedral is also discretized into dynamical states based on whether the dihedral is stable in a single rotameric state (ordered), or rapidly transitioning between multiple states (disordered) as described in earlier studies (Singh & Bowman, 2017; Sun et al., 2018).

From the probability distribution of the structural and dynamical states of every dihedral angle, a holistic communication for every pair of dihedrals is computed:

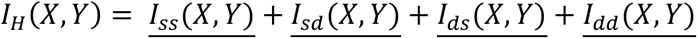

<u>*I*_*SS*_(*X*, *Y*)</u> is the normalized mutual information between the rotameric states of dihedral *X* and dihedral *Y*, <u>*I*_*Sd*_(*X*, *Y*)</u> is the normalized mutual information between the rotameric state of dihedral *X* and the dynamical state of dihedral *Y*, <u>*I*_*dS*_ (*X*, *Y*)</u> is the normalized mutual information between the dynamical state of dihedral *X* and the rotameric state of dihedral *Y*, and <u>*I*_*dd*_ (*X*, *Y*)</u> is the normalized mutual information between the dynamical state of dihedral *X* and the dynamical state of dihedral *Y*. The Mutual information (*I*) is computed as,

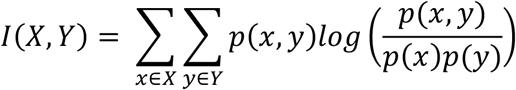

### Community detection

To detect clusters of residues strongly inter-communicating residues from the CARDS allosteric network, we applied Louvain community detection method (Blondel et al., 2008). Louvain algorithm identifies clusters by maximizing the modularity function defined as following (Fig. S2):

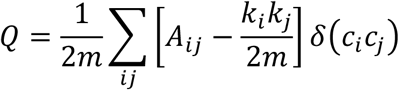

Here, *A*_*ij*_ is the edge-weight connecting two nodes *i* and *j*; *k*_*i*_ and *k*_*j*_ are the sum of all edge-weights connected to nodes *i* and *j*, respectively. *m* is the sum of all the edge-weights in the network. *c*_*i*_and *c*_*j*_ are the clusters of nodes *i* and *j*, respectively. *δ*=(*c*_*i*_*c*_*j*_) is a Kronecker delta function such that *δ*=(*c*_*i*_*c*_*j*_) =1 if two nodes *i* and *j* are in the same cluster, 0 otherwise.

### Plasmid construction

pBAD 6xHis-Gαi1 was a gift from Dr. Sergej Djuranovic, Washington University in St. Louis, St. Louis, MO (Verma et al., 2019).Further mutations were generated by site directed mutagenesis using specifically designed and synthesized primers for each mutation (Integrated DNA Technologies). All plasmids were verified by Sanger sequencing (GeneWiz).

### Protein expression and purification

All G proteins were expressed recombinantly in TOP10 cells (ThermoFisher). 0.5 L cultures were auto-induced for 72 h at room temperature in 2.4 L Erlenmeyer flasks with media containing 2% tryptone, 1% yeast extract, 1% glycerol, 0.5% NaCl, 50mM KH_2_PO_4_, and 25mM (NH_4_)_2_SO_4_ buffered to pH 8.0 and supplemented with 100μg/mL ampicillin, 2mM MgSO_4_, 0.4x Trace Metals, 5mM D-glucose, and 0.1mM L-arabinose at time of induction (“G ProteinsTechniques Anal.,” 1999; Studier, 2005). Induced cells were collected by centrifugation (4°C, 4,000 x g for 15 min), the supernatant was discarded, and the pellets were snap frozen in liquid nitrogen and stored at -80°C until purification. Harvested cells were thawed on ice, suspended in Lysis Buffer (50 mM Tris pH 8, 150 mM NaCl, 150 mM KCl, 25mM (NH_4_)_2_SO_4_, 10% glycerol, 10 mM BME, 1x cOmplete™ EDTA-free Protease Inhibitor Cocktail (Roche)), and then lysed by sonication on ice. The supernatant was cleared by centrifugation (100,000 x g for 30min) and then bound to Ni-NTA Agarose (Qiagen) for 1h at 4°C with shaking. Agarose resin was collected by centrifugation at 4°C (4,000 x g for 5min) and the supernatant discarded. Beads were washed twice with 50mL of Wash Buffer (50 mM Tris pH 8.0, 150 mM NaCl, 150 mM KCl, 25mM (NH_4_)_2_SO_4_, 10 mM BME, 10% glycerol) and shaking for 5 min at 4°C, transferred to a disposable gravity flow column and washed once with 25mL Wash Buffer. Protein was eluted with Elution Buffer (50 mM Tris pH 8.0, 150 mM NaCl, 150 mM KCl, 25 mM (NH_4_)_2_SO_4_, 10% glycerol, 500 mM imidazole) and a sample was taken for activity assays. The eluted protein was then dialyzed in Dialysis Buffer (50 mM HEPES, 10 mM MgCl_2_, 1 mM EDTA, 1 mM DTT, 10% glycerol, pH 8.0) starting with dialysis in 500 mL for at least 4 hours, then 500 mL overnight, and finally 1 L for at least 4 hours. Protein concentration was determined spectroscopically in Dialysis Buffer using an extinction coefficient of 35870 M^-1^ cm^-1^. Eluted protein was then aliquoted, snap frozen in liquid nitrogen, and stored at -80°C until use.

### Nucleotide exchange assays

Nucleotide loading activity of purified Gα subunits was detected with BODIPY-FL-GTPγS, a non-hydrolyzable fluorescent analog of GTP, and a Synergy H4 plate reader26. For association assays, BODIPY-FL-GTPγS (25 nM final concentration) was aliquoted into the wells of a black-wall clear-bottom 96-well plate (catalog number 3603, Costar). Nucleotide loading was initiated by injecting Gα subunits (1 μM final concentration) in Reaction Buffer (50 mM Tris-HCl pH 8.0, 1 mM EDTA, 10 mM MgCl_2_). For dissociation assays, BODIPY-FL-GTPγS (25 nM final concentration) and Gα subunits (1 μM final concentration) were preincubated for 1 hour in the wells of a black-wall clear-bottom 96-well plate (catalog number 3603, Costar) before injecting GDP (20 μM final concentration) in Reaction Buffer + 25 nM BODIPY-FL-GTPγS to initiate dissociation. Fluorescence intensity was monitored in well mode at 30°C for 15min with 1s intervals for association or in plate mode for 180 min with 5s intervals for dissociation with excitation and emission bandpass filters with wavelengths of 485/20 nm and 528/20 nm, respectively, and a 510 nm dichroic mirror. Resulting curves were fit to one-phase association models for k_obs_ and one-phase decay models for k_d_ using GraphPad Prism v6.01. From these values, k_a_ was determined from the relationship k_obs_ = [L] • k_a_ + k_d_ which, when rearranged, yields k_a_ = (k_obs_-k_d_)/[L]. Kd values were subsequently calculated using the relationship K_d_ = k_d_/k_a_.

## Results and discussion

### Automated identification of the allosteric network in Gq recapitulates prior results

To assess how well the allosteric network responsible for G protein activation is conserved, we first sought to develop an automated method for detecting allostery that would ensure robust and reproducible results. Many methods have been developed to dissect allosteric networks (Feher et al., 2014; Ichiye & Karplus, 1991; Lange & Grubmüller, 2006; Weinkam et al., 2012; Whitley & Lee, 2009). Most focus on concerted structural changes. For this work, we chose to use CARDS (**<u>C</u>**orrelation of **<u>A</u>**ll **<u>R</u>**otameric and **<u>D</u>**ynamical **<u>S</u>**tates, see supporting information for details, Fig. S1) (Singh & Bowman, 2017). CARDS is unique in its ability to account for coupling between both the structure and dynamics of pairs of dihedral angles. We previously showed that the dynamical insight it provides is important for understanding allostery in G proteins (X. Sun et al., 2018). While determining the coupling between pairs of dihedrals and coarse-graining this information to determine the coupling between pairs of residues is automated, further interpretation of the allosteric network previously required manual input. For instance, in our past work, we manually identified sets of residues with strong coupling to one another that spanned the GPCR- and nucleotide-binding sites (cite X. Sun et al). While this analysis provided significant insight, performing the same analysis on multiple isoforms is labor intensive and potentially introduces an unwanted element of subjectivity into any claims about conservation. Therefore, we sought to develop an automated process of identifying allosteric networks based on CARDS.

To automate the process of finding allosteric networks, we employed a community detection method called the Louvain algorithm which identifies clusters of strongly interconnected nodes within a network (Blondel et al., 2008). Like many clustering algorithms, the Louvain algorithm seeks to group together nodes (in this case, residues) that have strong links to each other while separating weakly connecting nodes into separate clusters. This methods has a number of desirable capabilities, particularly the ability to determine the number of clusters automatically. At the start of this method, each node in a network is assigned to its own cluster. In subsequent steps, these nodes are iteratively grouped together into larger clusters to maximize a quantity called modularity, which is a measure of the density and strength of connections among nodes within a cluster compared to connections between clusters.

Applying Louvain clustering to CARDS data from simulations of Gαq recapitulated insights from our past work, as expected (X. Sun et al., 2018). Louvain clustering identified six clusters of residues. We will refer to the three in the G domain as GDC1-3, the one in the H domain as HdC1, and the three at the interdomain interface IDC1-3 (Fig. 2). GDC1 encompasses the GPCR-binding interface (the C-terminal segment of H5 and the hNs1 loop), β strands S1-S4, residues from helix H1, and the P-loop that engages the nucleotide phosphates, implying that these regions have strong structural and dynamical correlations. GDC1 also has strong coupling to GDC3 (Fig. 2B), which includes other key components of the nucleotide-binding site, such as the switch-I and switch-II motifs which contain the active site residues responsible for GTP hydrolysis and undergo important nucleotide-dependent structural changes that distinguish the active and inactive states (Kamato et al., 2015; Van Eps et al., 2006; Wall et al., 1998). We refer to GDC1 and GDC3 as pathway-I for communication between the GPCR- and nucleotide-binding sites. GDC2 constitutes a second pathway, which we refer to as pathway-II, as it encompasses GPCR-binding elements like β strand S6 and nucleotide-binding elements like helix HG and the s6h5 loop. These pathways are consistent with previous computational studies of allostery in Gq and other G protein isoforms (Dror et al., 2015; Flock et al., 2015; Sun et al., 2018).

**Figure 2.**
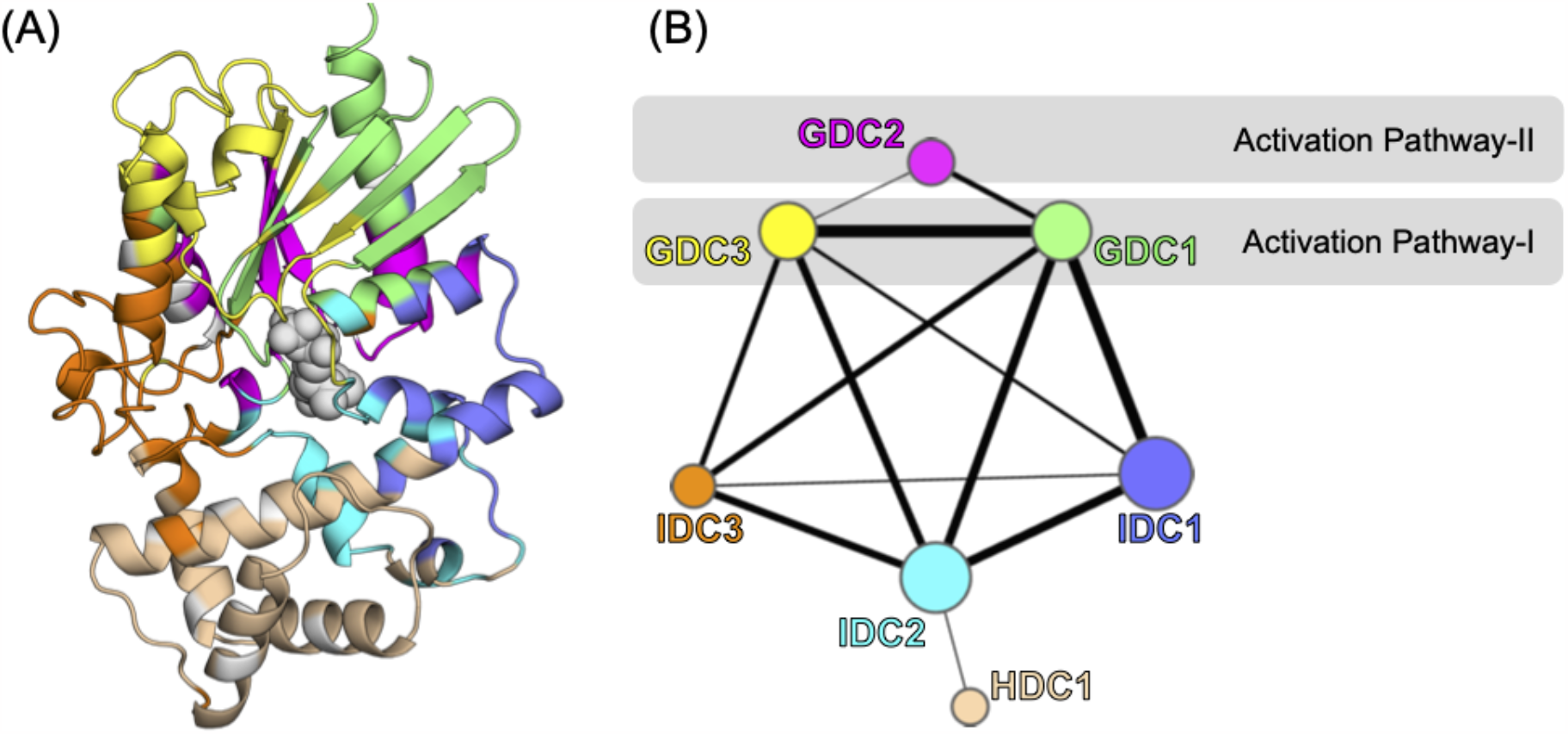
The allosteric network in Gq that was automatically identified by Louvain clustering. (A) A structure of Gq with residues colored based on which of the six clusters they are grouped into. (B) A network representation of the allosteric network showing the color coding of the G domain clusters (GDC1-3), the H domain cluster (HDC1), and the interdomain clusters (IDC1-3). Each node represents a cluster, the size of the node is proportional to the average strength of CARDS communication between amino acids within the cluster, and the width of the edges between nodes is proportional to the average strength of CARDS communication between residues in the two clusters.

### Automated analysis of other isoforms suggests similar allosteric networks

Encouraged by the success of Louvain clustering in finding important allosteric pathways in Gq, we extended this approach to other G*α* isoforms to understand and compare their allosteric dynamics. We selected at least one representative isoform from each of the other families besides Gq/11, specifically Gi1, Gt, and Go from the Gi family, G13 from the G12/13 family, and Gs from the Gs family. We collected ∼35 microseconds of molecular dynamics simulation data for each isoform and applied CARDS to quantify the coupling between every pair of dihedral angles. Then we applied Louvain clustering to coarse-grain these networks and begin understanding how similar they are.

Examining the allosteric networks identified by Louvain clustering suggested that the different isoforms share many common features. For example, several residues from the C-terminal of helix H5, β strands S1-S3 and helix H1 are in a common cluster in multiple isoforms (Fig. 3), suggesting these regions are strongly coupled across subfamilies. These regions are an important part of pathway-I, suggesting this path may be shared by many isoforms. Similarly, parts of activation pathway-II, such as residues from β strand S6 and the s6h5 loop, are grouped together in all the isoforms. There are also similarities outside the activation pathways. For example, a large part of the H domain is grouped into a single cluster in multiple isoforms.

**Figure 3.**
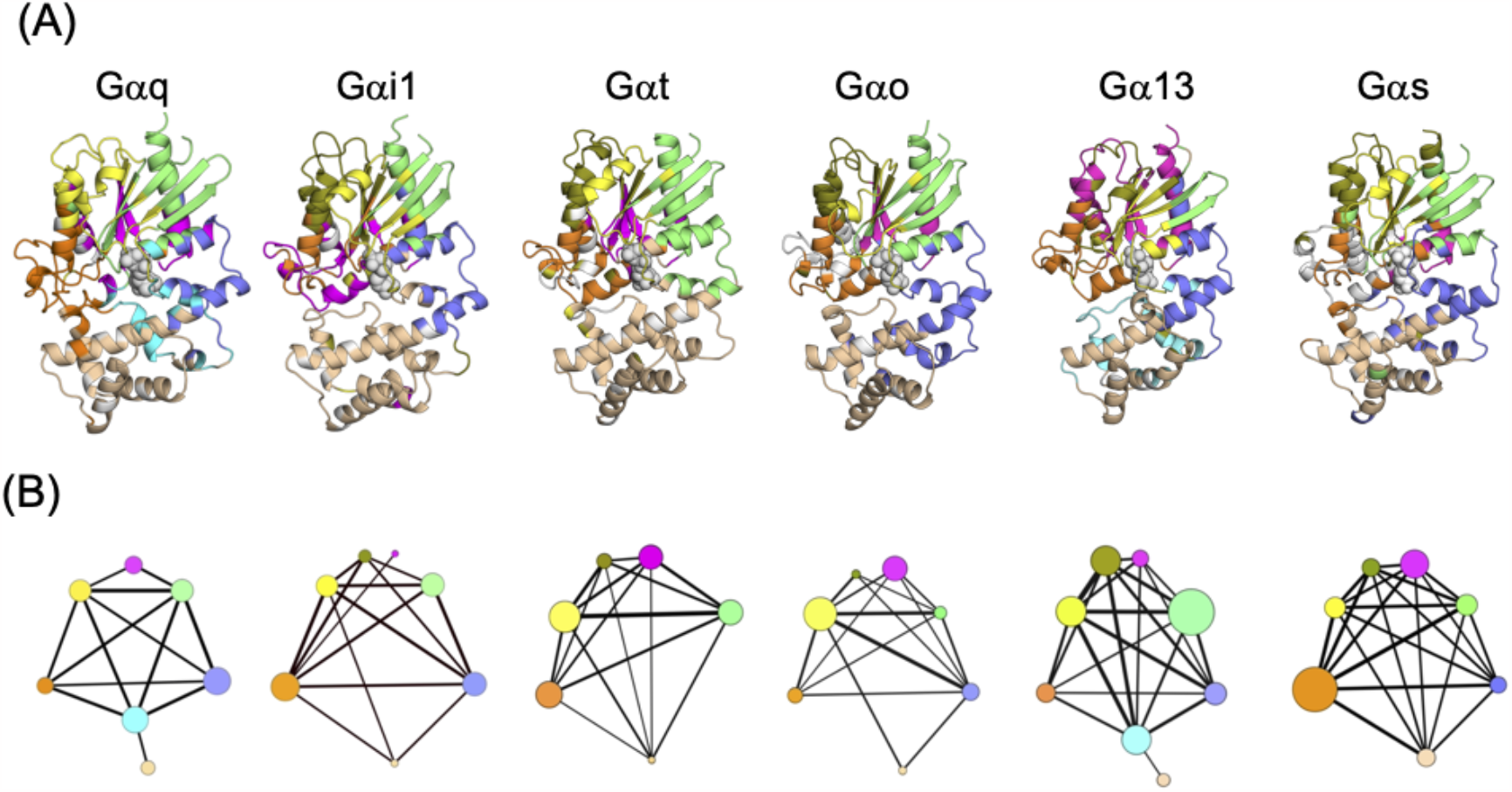
Allosteric networks for six different G proteins covering all four subfamilies. (A) Louvain clustering of CARDS communication networks for G protein isoforms (Gq, Gi1, Gt, Go, G13, and Gs) mapped on their structures. (B) Network diagrams showing coupling between clusters in each G protein isoform shown in (A). Each node is a cluster with a size proportional to the average strength of coupling between residues in that cluster. Edge widths are proportional to the strength of communication between clusters.

Despite the common allosteric aspects discussed above, we observed significant differences in the number and composition of clusters among in the different isoforms that could be useful for targeting specific subsets of them. For example, the residues in GDC2 of Gq are split between two clusters in Gt, suggesting that pathway-II may be weaker in Gt than it is in Gq. Parts of switch-II also were assigned to different clusters in different isoforms. These differences are noteworthy because switch-II is important for nucleotide binding and is part of the binding interface that G proteins use to engage effector molecules. Therefore, differences between the allosteric coupling that controls the conformation of switch-II in various isoforms could have important functional consequences.

### Quantifying the conservation of allostery between isoforms identifies a universal activation mechanism

While the similarity between the allosteric networks identified by Louvain clustering is suggestive of a large degree of conservation, we employed further analyses to establish whether the allosteric networks of the different Gα isoforms are similar at the residue level.

To quantify the extent of conservation at the residue level, we adapted a network alignment method originally developed to study how protein-protein interaction networks are conserved between organisms (Berg & Lässig, 2006). The first step was to create an alignment network from two parent networks (A and B) one wishes to compare. Each node in the alignment network represents a pair of equivalent nodes from the parent networks. In this case, a node in the alignment network refers to equivalent nodes from two Gα isoforms, where equivalency is determined by the CGN numbering scheme. Edges between nodes in the alignment network quantify how strongly conserved the allosteric coupling between a pair of residues is. This conservation score (∆*s*(*a, b*)) is defined as

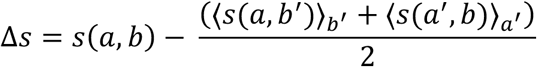

Where *a* is the strength of coupling between the two residues in parent network A and *b* is the strength of coupling between the two residues in parent network B. 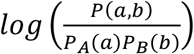, *P*(*a, b*) is the joint probability that allosteric correlations between equivalent residues in two isoforms A and B have strengths *a* and *b*, respectively, *P*_*A*_(*a*) is the probability that an allosteric correlation has strengths *a* in the isoform A. ⟨*s*(*a, b*^′^)⟩_*b′*_ is the average over all aligned inter-residue allosteric correlations with strength *a* fixed, and ⟨*s*(*a*^′^, *b*)⟩_*a′*_ is the average over all aligned inter-residue allosteric correlations with strength *b* fixed. We used Gq as a reference, and compared each isoform to it. Further details are given in the supporting information.

This approach confirmed that the different Gα isoforms share a largely conserved allosteric network and highlighted specific residues of importance (Figs. 4 and 5). Many of the inter-residue correlations that constitute activation pathway-I in Gq are conserved in multiple isoforms. For example, allosteric coupling between residue Phe^G.H5.8^ from helix H5, several residues from β strands S1-S3, Lys^G.H1.1^, Met^G.H1.8^, Ile^G.H1.11^, and His^G.H1.12^ from helix H1 and the P-loop were found to be conserved in all isoforms. Similarly, key parts of activation pathway-II are conserved, such as Tyr^G.S6.2^, His^G.S6.4^ and Phe^G.S6.5^ from β strand S6 that engages GPCRs and residues Thr^G.s6h5.1^ and Cys^G.s6h5.2^ from the s6h5 loop that engages the nucleotide. The binding site of a known Gq inhibitor, called YM-254890 (YM), is also part of this conserved allosteric network, suggesting that molecules like YM might be designed to inhibit other Gα isoforms by targeting this region. This largely universal allosteric network is depicted in Fig. 5. Interestingly, the conservation of allosteric coupling does not necessitate the conservation of the sequence. Several of the key residues just highlighted are of different amino acid types in different isoforms.

**Figure 4.**
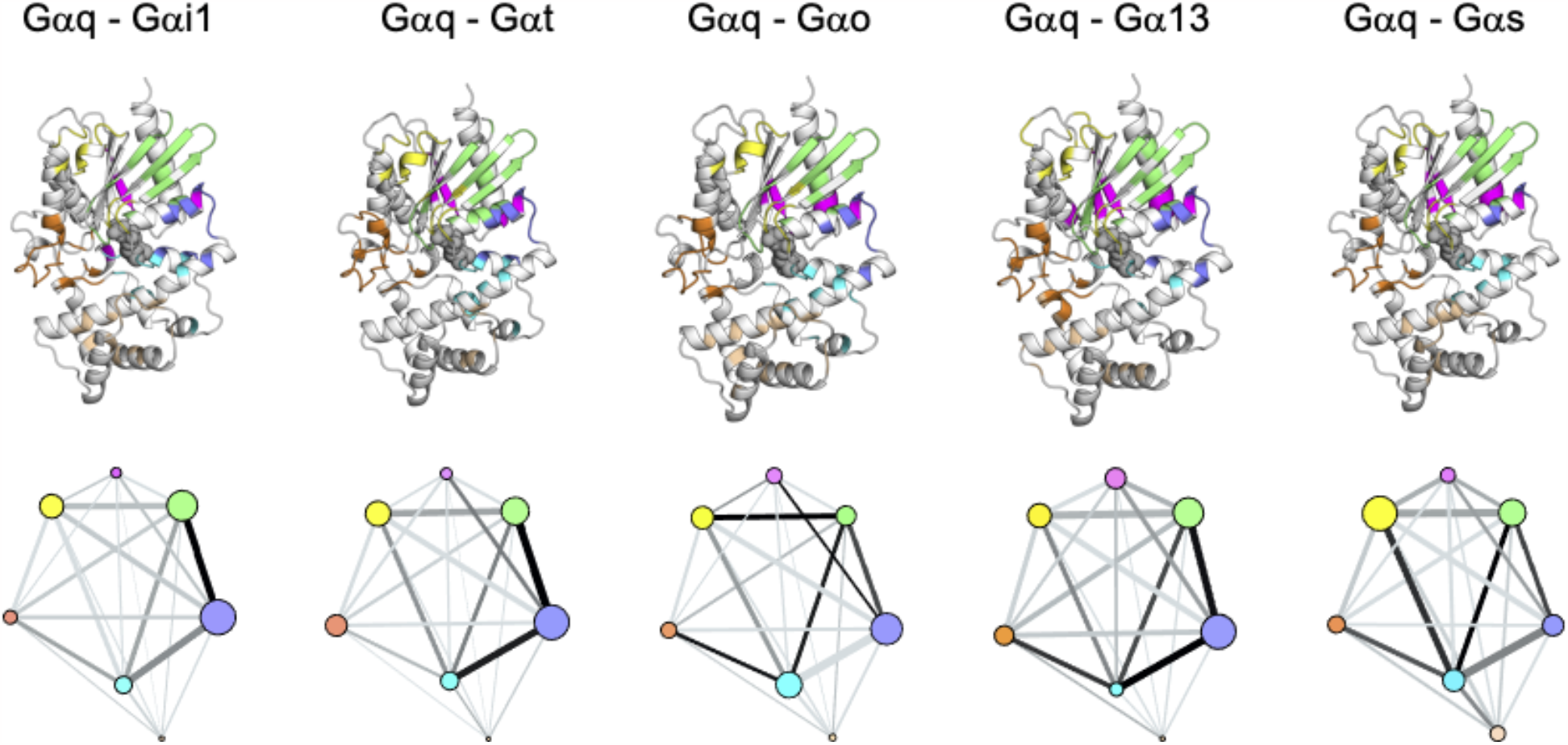
Conservation of the allosteric networks of each Gα isoform compared to Gq. The upper panel shows a representative structure with residues whose allosteric coupling is conserved color coded according to the network representation of the conserved allosteric network in the lower panel. Each node represents a group of amino acids with conserved allosteric coupling compared to Gq. Residues and nodes are colored based on the allosteric network identified for Gq. The size of each node is proportional the strength of coupling within that cluster for the given isoform and the edge width is proportional to the strength of coupling between clusters. The edge color is proportional to the conservation score, taken with respect to Gq (dark means highly conserved, light means weakly conserved).

**Figure 5.**
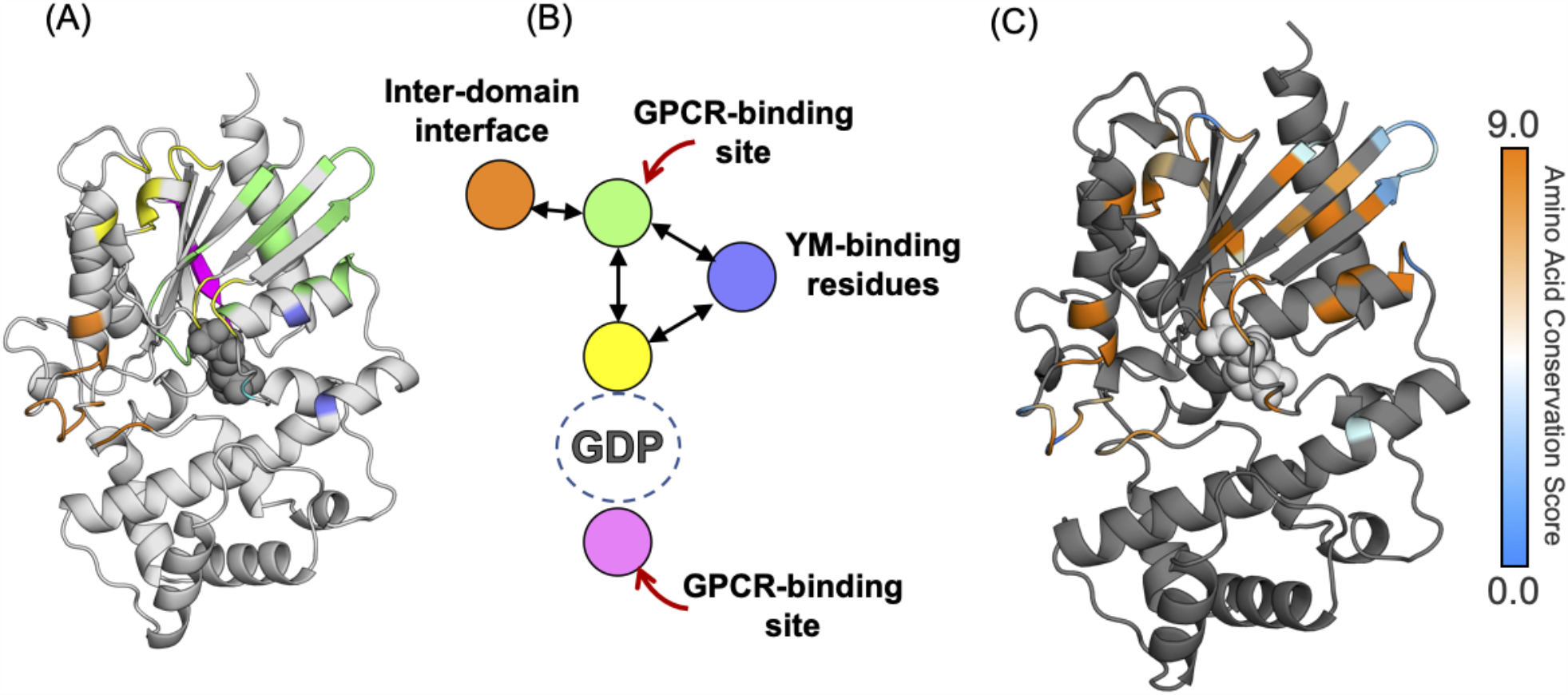
A largely universal allosteric network is not directly linked to sequence conservation. (A) Key residues mapped onto a representative structure and colored according to the allosteric network shown in (B). (C) Amino-acid conservation scores of residues in the unified allosteric network mapped on a representative structure. The amino-acid conservation score is computed on the Consurf server using a multiple sequence alignment of 16 G*α* isoforms.

### Activation pathway-I is the dominant communication channel

Examining the allosteric networks for each isoform and the conservation between them suggested that allosteric pathway-I (GDC1 and GDC3) is generally stronger than pathway-II (GDC2). Each of the clusters in pathway-I has stronger coupling between residues within the cluster than is found in pathway-II, as denoted by the larger sizes of GDC1/3 compared to GDC2 in the network diagrams (Figs. 3 and 4). The coupling between GDC1 and GDC3 is typically stronger than the coupling of GDC2 to any other cluster, again supporting the greater strength of pathway-I. Within pathway-I, our analysis of the conservation of allosteric coupling suggests that Phe^G.H5.8^ has strongly conserved coupling with the S1-S3 strands and helix H1, notably Phe^G.S2.6^, Phe^G.S3.3^, Met^G.H1.8^ and His^G.H1.12^ (Figure 4 & Figure 5A). The strength of this coupling is consistent with analysis of amino acid conservation by Flock et al., who highlighted the hydrophobic interactions of Phe^G.H5.8^ with Phe^G.S2.6^ and Phe^G.S3.3^ from the S1-S3 strands, as well as Met^G.H1.8^ and His^G.H1.12^ from the H1 helix (Flock et al., 2015). Breaking the allosteric coupling we observe down into contributions from the side-chains and the backbone suggests that the side-chains are more important (Fig. S3). Therefore, mutating Phe^G.H5.8^ is likely to disrupt pathway-I. This prediction is also consistent with a previous simulation study (Louet et al., 2012). Similarly, His^G.S6.4^ stood out as having strong and conserved coupling within pathway-II. The side-chain of this residue also appears to be more important than the backbone (Fig. S4), suggesting mutating His^G.S6.4^ is likely to disrupt pathway-II.

To test this hypothesis, we mutated the key residues identified in each pathway and experimentally assessed their effect on nucleotide exchange. As in previous work (Kaya et al., 2014; Marin et al., 2001), we reasoned that mutations that increase the basal rate of nucleotide exchange are disrupting allostery. We focused on Gi because it has the biggest difference between the strength of pathway-I and pathway-II (Fig. 3).

Our results from mutagenesis and nucleotide exchange experiments corroborated the prediction that pathway-I is stronger than pathway-II (Fig. 6 and Table 1). We found that introducing the mutation Phe336^G.H5.8^Leu to disrupt pathway-I in Gi resulted in a greater than 2-fold increase in nucleotide exchange rate. Previous work also showed that the equivalent mutation in Gt results in enhanced nucleotide exchange (Marin et al., 2001), consistent with our prediction that this allosteric pathway is conserved across multiple Gα isoforms. Furthermore, the equivalent substitution in Ras has also been observed to accelerate nucleotide exchange (Quilliam et al., 1995), suggesting that the conservation of pathway-I extends beyond Gα subunits of heterotrimeric G proteins to the large family of monomeric G proteins. Meanwhile, introducing the His322^G.S6.4^Ala mutation to disrupt pathway-II did not have a statistically significant effect on nucleotide exchange rate. This result is also consistent with a previous study that found that Tyr320^G.S6.2^Ala in pathway-II had little effect (D. Sun et al., 2015). To further test our hypothesis, we selectee two additional mutations to pathway-I, Met53^G.H1.8^Leu and His^G.H1.12^Phe. Both mutations resulted in enhanced nucleotide exchange, again consistent with the importance of pathway-I.

**Table 1.** BODIPY-FL-GTPγS exchange rates for Gαi1 WT and mutants.

| Protein | $k_{obs}$<br>(min<sup>-1</sup>) | $k_d$<br>(min<sup>-1</sup>) | $k_a$<br>(M<sup>-1</sup>.min<sup>-1</sup>) | $K_D$<br>(M) |
| --- | --- | --- | --- | --- |
| WT | 0.17 | 0.031 | 5.6E+06 | 5.7E-09 |
| Met53Leu | 1.2 | 0.35 | 3.2E+07 | 1.1E-10 |
| His57Phe | 0.28 | 0.073 | 8.2E+06 | 8.9E-09 |
| His322Ala | 0.15 | 0.055 | 3.7E+06 | 1.5E-10 |
| Phe336Leu | 0.43 | 0.23 | 8.0E+06 | 2.9E-10 |

**Figure 6.**
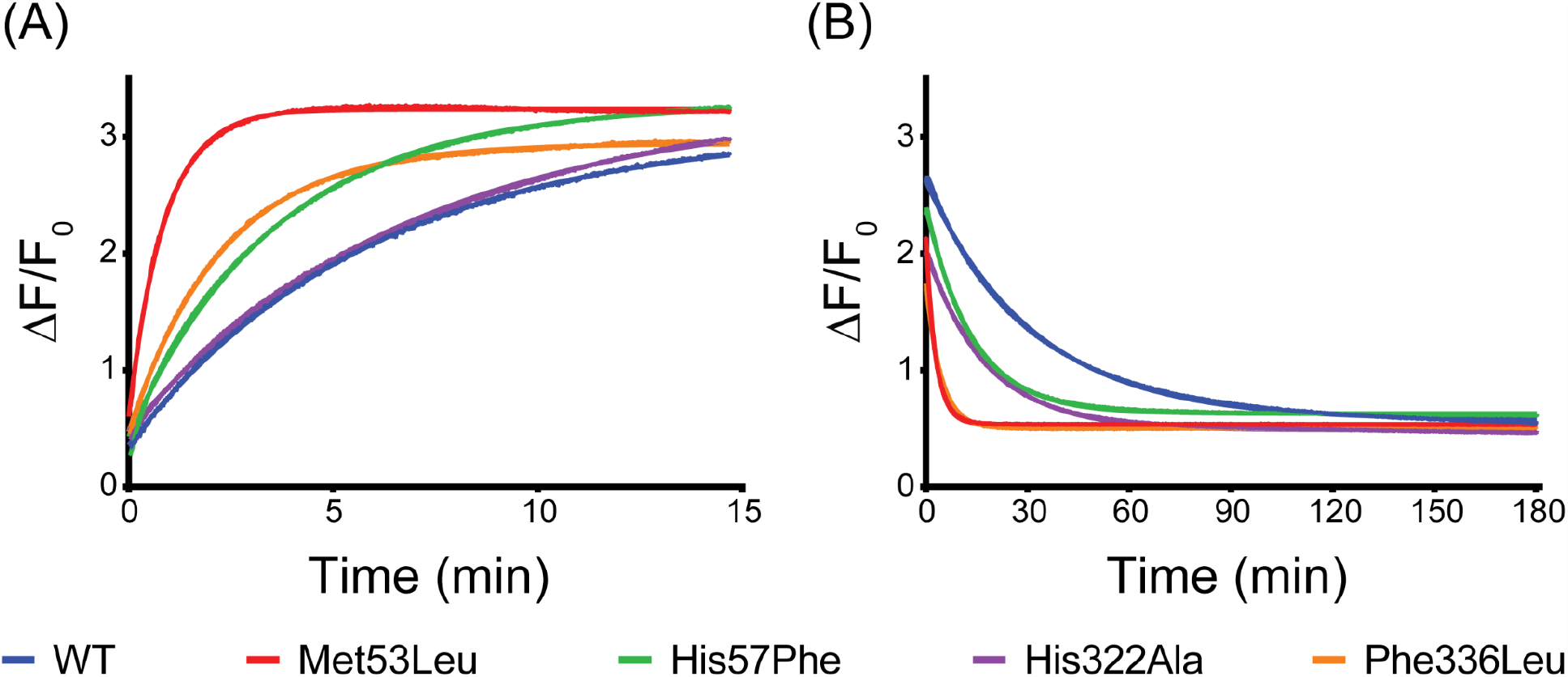
Mutations in pathway-I have a significant impact on nucleotide exchange while mutations in pathway-II do not. (A) BODIPY-FL-GTPγS loading of each mutant. Curves are the average of 9 independent measurements with fitted curves. (B) BODIPY-FL-GTPγS dissociation of each mutant. Curves are the average of 6 independent measurements with fitted curves.

## Conclusions

We have identified a largely universal mechanism of G protein activation by directly observing allosteric coupling in molecular dynamics simulations of representatives of all four subclasses of Gα subunits. Our results are largely consistent with previous inferences from sequence conservation (Flock et al., 2015). This complementarity is important as our ability to see allostery in action is free of confounding factors based purely on sequence conservation, such as requirements for protein stability. Our models also provide insight into elements of the allosteric coupling that are unique to different isoforms and, therefore, may help inform the design of isoform-specific drugs. The automated tools for inferring allostery and conservation that we have developed suggest there are two dominant pathways for allosteric coupling in G proteins, which we call pathway-I and pathway-II. Our models suggest that pathway-I is dominant, and this finding is supported by experimental characterization of mutations designed to disrupt the two pathways. We expect our contributions to understanding G protein biology will help inform the design of new inhibitors. More broadly, our tools should also be useful for studying allostery in other proteins and assessing the extent to which this allostery is conserved in related proteins.

## Acknowledgements

We are grateful to the Folding@home community for volunteering their personal computing power to run simulations for this work. This work was funded by NIH R01 GM124093 (to K.J.B.) and GM124007, as well as NSF MCB-2218156. G.R.B. holds a

Packard Fellowship from the David and Lucile Packard Foundation.

