## Supplementary material for "G protein activation occurs via a largely universal mechanism": Supporting Information.pdf

**Fig. S1**

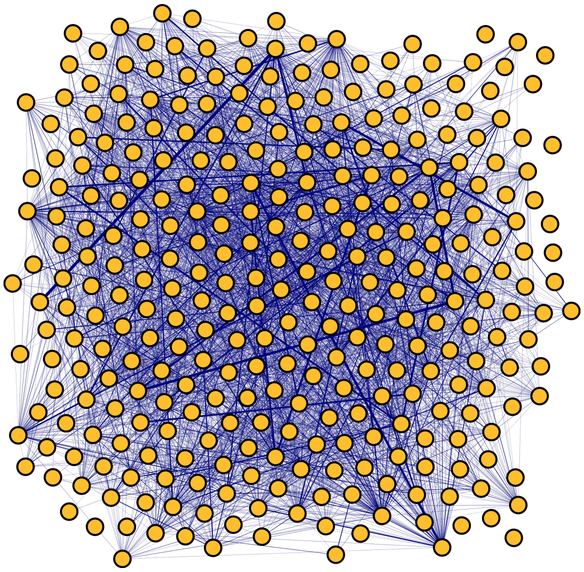

**Figure S1.** An illustrative diagram showing inter-residue CARDS network. Each yellow sphere (node) represents a residue and the blue line (edge) connecting two nodes corresponds to inter-residue allosteric communication. Width of the edge is proportional to the strength of allosteric communication.

**Fig. S2**

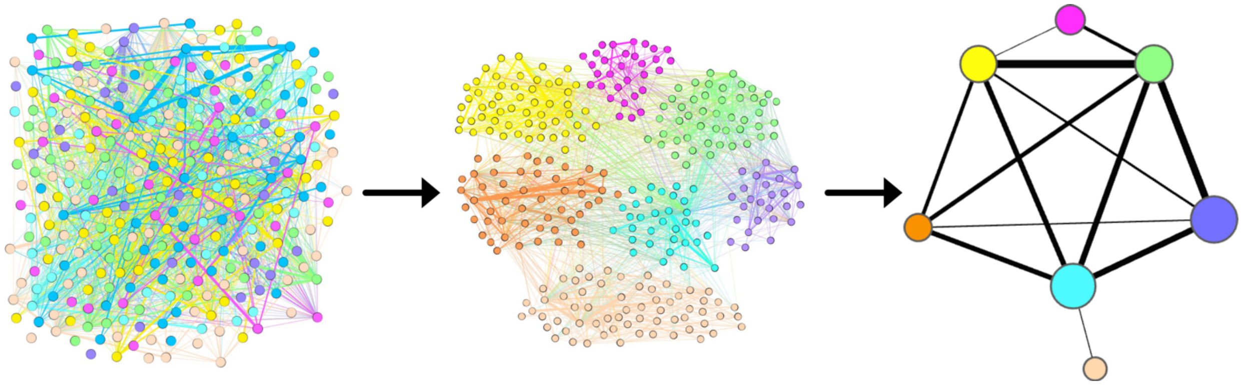

**Figure S2.** An illustrative diagram depicting Louvain clustering. The network shown on the left is an inter-residue CARDS communication network. Each node (residue) in this network is colored by the Louvain cluster it belongs to. The edge connecting two nodes corresponds to inter-residue allosteric communication. The network shown in the middle shows the rearrangement of the nodes based on modularity. Louvain clusters consisting of residues with strong allosteric communications among themselves are shown in the network shown on the right. The edge connecting two Louvain clusters (nodes) corresponds to the strength of average allosteric communication between the two clusters.

**Figure S3**

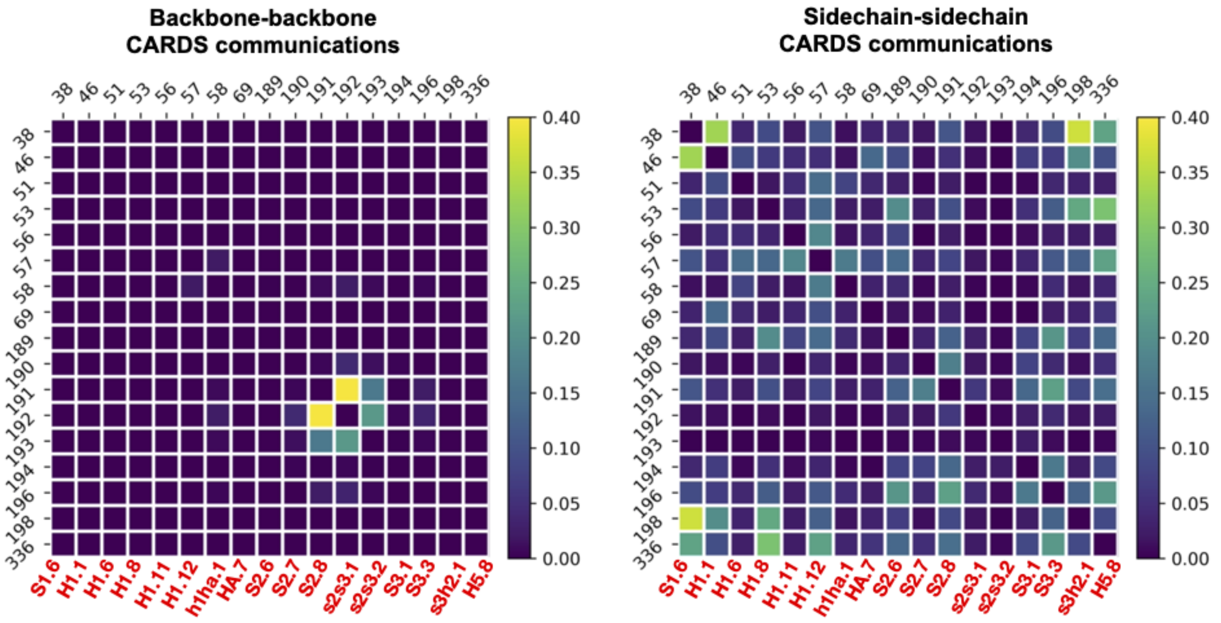


**Figure S3.** (A) Backbone-backbone CARDS communications in conserved allosteric activation pathway-I. (B) Sidechain-sidechain CARDS communications in conserved allosteric activation pathway-I. Axis labels are residue numberings in Gi1 isoform. CGN nomenclature for each residue is denoted in red font.

**Figure S4**

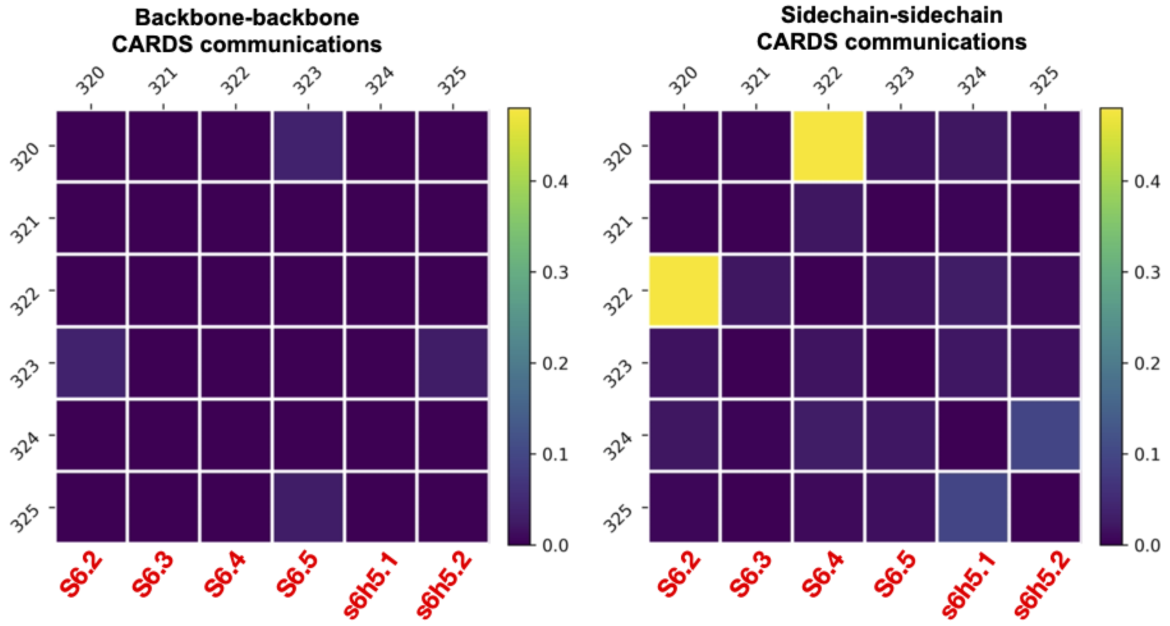


**Figure S4.** (A) Backbone-backbone CARDS communications in conserved allosteric activation pathway-II. (B) Sidechain-sidechain CARDS communications in conserved allosteric activation pathway-II. Axis labels are residue numberings in Gi1 isoform. CGN nomenclature for each residue is denoted in red font.
